# Exploring the effects of metformin on the body by urine proteome

**DOI:** 10.1101/2024.12.30.630710

**Authors:** Yuzhen Chen, Haitong Wang, Minhui Yang, Ziyun Shen, Youhe Gao

**Author notes:** Corresponding author: Prof. Youhe Gao;.

## Abstract

Metformin is the first-line medication for treating type 2 diabetes mellitus (T2DM), with over 200 million patients taking it daily. Its effects are extensive and play a positive role in multiple areas. Can we explore its effects and potential mechanisms by urine proteome? In this study, a total of 166 differential proteins were identified after rats were given a dose of 150 mg/(kg·d) of metformin for 5 consecutive days, including complement component C6, pyruvate kinase, coagulation factor X, growth differentiation factor 15 (GDF15), carboxypeptidase A4, chymotrypsin-like elastase family member 1, and L-lactate dehydrogenase C chain (LDH-C). Several of these proteins have been reported to be directly affected by metformin or associated with the effects of metformin. Several biological pathways enriched by the differential proteins or proteins where the differentially modified peptides are located have been reported to be associated with metformin, including glutathione metabolic process, negative regulation of gluconeogenesis, and renin-angiotensin system. Additionally, some significantly enriched biological pathways that have not been reported to be related to the effects of metformin may provide clues for the study of metformin’s potential mechanisms. In conclusion, the application of urine proteome offers a comprehensive and systematic approach to exploring both the known and unknown effects of drugs, thus opening a new window to study the mechanisms of metformin.

## 1 Introduction

Metformin, the first-line medication for treating type 2 diabetes mellitus (T2DM) in clinical use for over 60 years, is taken daily by more than 200 million T2DM patients globally [1]. In addition to lowering blood glucose, it also plays a positive role in cognitive function improvement [2], tumor suppression [3], cardiovascular protection [4], antiaging [5], and weight loss [6]. Metformin shows good safety and tolerability in T2DM treatment [7], with no increased hypoglycemia risk when used alone [8]. Metformin is also cost-effective compared to other hypoglycemic drugs and has been recommended by several guidelines as a basic treatment for controlling hyperglycemia in patients with T2DM [1,9].

In recent years, many researchers have continued exploring metformin’s effects and potential mechanisms. Metformin was found to promote the biosynthesis of *N*-lactoyl-phenylalanine (Lac-Phe), suppressing appetite and leading to weight loss [10,11]. By increasing the abundance of *Akkermansia muciniphila* in the gut, it regulates the pathways associated with inflammation in the host body, reduces the level of proinflammatory cytokines in plasma, and improves cognitive function [12]. Metformin may reprogram tryptophan metabolism and drive immune-mediated antitumor effects [13]. It can also activate Nrf2, a transcription factor with antioxidant ability, and delay the aging of neurons and brain [14]. Although metformin has been widely used, its mechanisms have not been fully elucidated.

With the development of high-throughput sequencing technology, proteomics research has been deepened, which can reveal the composition and changing law of proteins in cells or organisms by analyzing protein structure, expression, post-translational modifications, and protein interactions [15]. Urine, which is not strictly regulated by homeostatic mechanisms, can accommodate and accumulate more changes, reflecting changes in all organs and systems of the body earlier and more sensitively [16]. In addition, urinary proteins do not directly originate from drugs, and the changes in the urine proteome reflect the changes in the body under the influence of drugs. Therefore, the study using urine proteome can comprehensively and systematically reflect the overall effects of drugs on the body.

The urine proteome is inevitably affected by a variety of factors such as age [17], genetics [18], gender [19], diet [20], and exercise [21]. Thus, minimizing the influencing factors in experiments is important. Animal models, whose genetic and environmental factors can be controlled, are suitable choices [22].

Several studies have been conducted to show that the effects of drugs can be reflected in the urine proteome. For example, in a previous study, the effects of prazosin, an alpha1-adrenergic receptor antagonist, on the urine proteome of rats were explored. A total of 775 proteins were identified, approximately half of which were related to prazosin treatment [23]. Another study using anticoagulants to affect the coagulation status in rats showed enrichment of the coagulation system, intrinsic prothrombin activation pathway, and extrinsic prothrombin activation pathway. And even coagulation changes induced by drugs exerting different coagulation mechanisms can be distinguished by urine proteome [24].

In addition, post-translational modification (PTM) of proteins plays an important role in the regulation of protein function, which can change the chemical nature, structure, or function of proteins, and then affect their activity, localization, folding, interaction with other proteins, and regulate many physiological and pathological processes [25]. These are essential for increasing the diversity of the proteome and maintaining cellular homeostasis. Drug-regulated post-translational modifications of proteins can be used as molecular target binding markers to identify pathways of drug regulation, which can help to reveal drug targets and mechanisms of action [26]. However, the effects of drugs on proteomic post-translational modifications have received little attention.

Therefore, for metformin, a drug with a wide range of effects and many users, can we take advantage of the fact that urine can reflect the body’s state comprehensively, systematically, and sensitively to explore the effects and potential mechanisms of metformin meticulously? In this study, rat models were established by intragastric administration of metformin to explore the effects of metformin on the body through urine proteome. Exploring the known or unknown mechanisms of action of metformin opens a new window for the study of its mechanisms (as shown in Figure 1).

**Figure 1.**
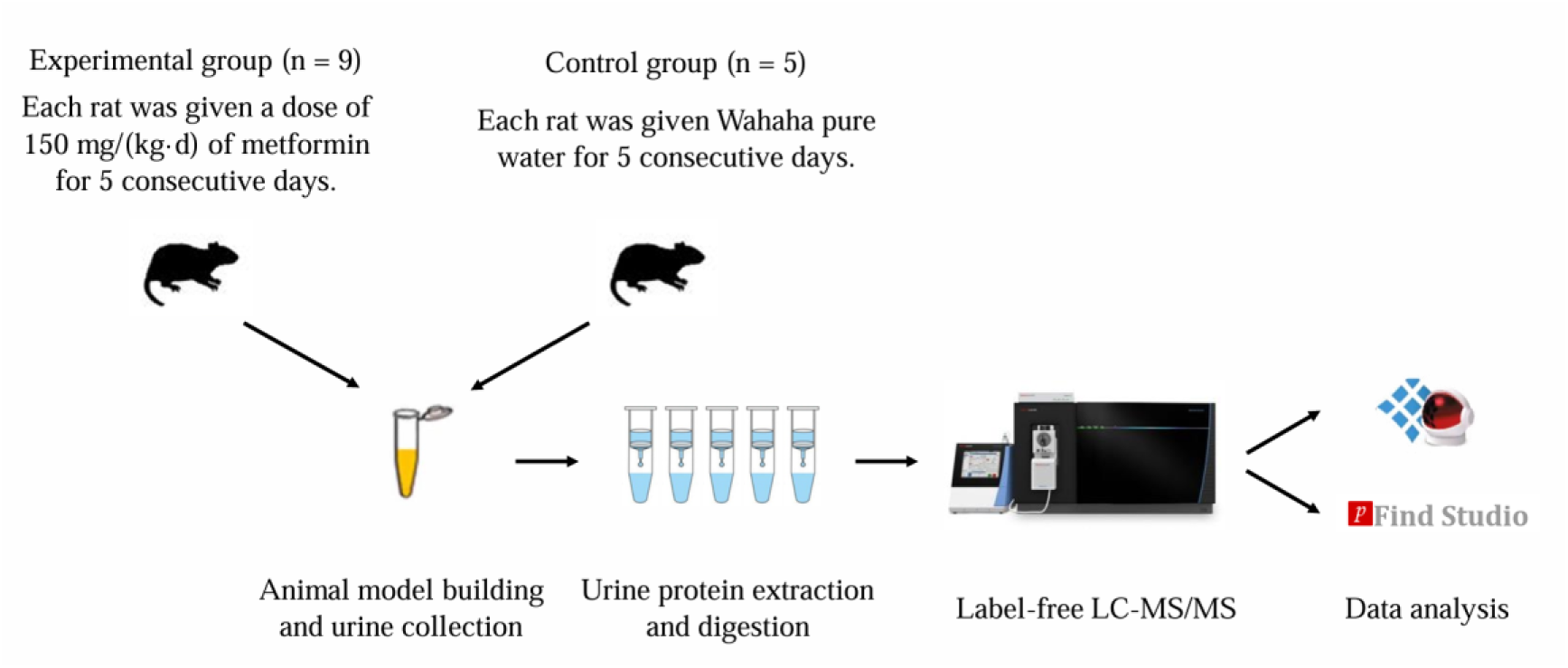
Technical route to explore the effect of metformin on the rat urine proteome.

## 2 Materials and Methods

### 2.1 Urine collection

14 healthy male Sprague Dawley(SD) rats (200±20 g) aged 6-7 weeks were purchased from Beijing Vital River Laboratory Animal Technology Co., Ltd. The experiment began after the rats were kept in a standard environment [room temperature (22±1) ℃, humidity 65%-70%] for 3 weeks when the rats weighed 370±30 g. The animal experiments were reviewed and approved by the Ethics Committee of the College of Life Sciences, Beijing Normal University (No. CLS-EAW-2020-034).

Metformin was dissolved in Wahaha pure water and administered intragastric once a day at the same time for 5 days. The experimental group (n=9) was given a dose of 150 mg/kg of metformin, which has been reported to achieve plasma concentrations of metformin in rats similar to those in humans [27]. The control group (n=5) was given Wahaha pure water. After 5 days of intragastric administration, rats were uniformly placed in metabolic cages, fasted, and water deprived, to collect urine samples for 12 h. The urine samples were stored at−80°C.

### 2.2 Urine sample preparation for label-free analysis

The collected urine samples were centrifuged at 12 000 ×*g* for 40 min at 4℃. The supernatant was transferred to a new centrifuge tube. Pre-cooled anhydrous ethanol with a volume three times that of the supernatant was added, which was homogeneously mixed, and then precipitated overnight at−20℃. The mixture was centrifuged at 12 000 ×*g* for 30 min at 4℃ and the supernatant was discarded. The protein precipitate was then suspended in an appropriate lysis buffer (8 mol/L urea, 2 mol/L thiourea, 25 mmol/L dithiothreitol, and 50 mmol/L Tris). After fully dissolved, the sample was centrifuged at 12 000 ×*g* for 30 min at 4℃. The supernatant was transferred to the new centrifuge tube to obtain urine protein extract. Protein concentration was quantified using the Bradford kit assay.

A total of 100 μg of protein was added into a 1.5 mL centrifuge tube. 25 mmol/L NH_4_HCO_3_ solution was added to make a total volume of 200 μL. 20 mmol/L dithiothreitol solution (DTT, Sigma) was added, vortexed, and mixed well, and heated in a metal bath at 97℃ for 10 min. After cooling to room temperature, 50 mmol/L iodoacetamide solution (IAA, Sigma) was added, vortexed, and mixed well, and reacted at room temperature without light for 40 min. 200 μL of UA solution (8 mol/L urea, 0.1 mol/L Tris-HCl, pH 8.5) was added to a 10 kD ultrafiltration tube (Pall, Port Washington, NY, USA) and centrifuged twice at 14 000 ×*g* for 5 min at 18℃. The treated protein sample was added and centrifuged at 14 000 ×*g* for 40 min at 18℃. 200 μL of UA solution was then added and centrifuged at 14 000 ×*g* for 40 min at 18℃, repeated once. 25 mmol/L NH_4_HCO_3_ solution was added and centrifuged at 14 000 ×*g* for 40 min at 18℃, repeated once. The samples were digested overnight at 37 ℃ with trypsin (Trypsin Gold, Promega, USA) (enzyme-to-protein ratio of 1:50). The digested peptides were eluted from ultrafiltration membranes, desalted with HLB columns (Waters, Milford, MA), dried in a vacuum desiccator, and stored at −80℃.

### 2.3 Liquid chromatography coupled with tandem mass spectrometry analysis

The digested peptides were dissolved in 0.1% formic acid, and the peptide concentration was quantified using a BCA kit. Then the peptides were diluted to a concentration of 0.5 μg/μL. A mixed peptide sample was prepared with 14 μL of each sample and separated using a high pH reversed-phase peptide fractionation kit (Thermo Fisher Scientific). Ten fractions were collected by centrifugation, dried using a vacuum desiccator, and redissolved in 0.1% formic acid. The indexed retention time (iRT) reagent (Biognosis) was then added (iRT-to-sample ratio of 1:10). For analysis, 1 μg of the peptide from each sample was loaded and separated using EASY-nLC1200 system with mobile phase A (0.1% formic acid) and mobile phase B (0.1% formic acid in 80% acetonitrile), eluted with a 90-min gradient as follows: 0 min, 4% phase B; 0 min–2 min, 6% phase B; 2 min–62 min, 22% phase B; 62 min–78 min, 35% phase B; 78 min–90 min, 90% phase B, and then analyzed with an Orbitrap Fusion Lumos Tribrid Mass Spectrometer (Thermo Fisher Scientific).

### 2.4 Database searching and data processing

To generate a spectral library, ten fractions obtained by centrifugation were analyzed with mass spectrometry in data-dependent acquisition (DDA) mode. The DDA results were then imported into Proteome Discoverer software to search against the *Rattus norvegicus* database with SEQUEST HT. The search results were used to establish the data-independent acquisition (DIA) method. The width and number of windows were calculated based on the *m/z* distribution density. Individual samples were analyzed using DIA mode. The pooled peptides were used to control the quality of the whole analytical process. The results were then imported into Spectronaut Pulsar software for analysis and processing. The peptide intensity was calculated by summing the peak areas of the respective fragment ions for MS^2^. The protein intensity was calculated by summing the respective peptide intensities. All results were then filtered according to a *Q* value less than 0.01 [corresponding to false discovery rate (FDR) less than 1%], and each protein contains at least 2 specific peptides.

The modification information of the proteome was obtained by pFind Studio software (Version 3.2.0, Institute of Computing Technology, Chinese Academy of Sciences, China). Data collected by the Orbitrap Fusion Lumos Tribrid Mass Spectrometer were analyzed for label-free quantification, and the original data files were searched against the *Rattus norvegicus* UniProt canonical database (updated to September 2024). For the search, “HCD-FTMS” was selected for “MS Instrument”, “Trypsin_P KR P C” was selected for trypsin digestion, and the maximum number of missed cleavages allowed per peptide was two. Both precursor tolerance and fragment tolerance were set as ±20 ppm. To discover global modifications, “Open Search” was selected. The FDRs at spectra, peptide, and protein levels were less than 1%, and the *Q* value at the protein level was less than 1%. The number of peptide mass spectra of each sample was extracted from the analysis results of pFind Studio using the script “pFind_protein_contrast_script.py” [28,29]. The modified peptides with reproducibility > 50% were screened.

### 2.5 Data analysis

The proteins and the number of the modified peptide mass spectra identified in the experimental and control groups were compared separately. The differential proteins and differentially modified peptides were screened with the following criteria: fold change (FC) ≥1.5 or ≤ 0.67, and *P* < 0.05 by two-tailed unpaired *t*-test analysis.

Hierarchical cluster analysis (HCA) and principal component analysis (PCA) were performed using the SRplot web server (http://www.bioinformatics.com.cn/). Functional enrichment of differential proteins was performed using the Database for Annotation, Visualization, and Integrated Discovery (DAVID). The reported literature was searched in the PubMed database (https://pubmed.ncbi.nlm.nih.gov) to conduct the functional analysis of the differential proteins and proteins where the differentially modified peptides are located.

## 3 Results and Discussion

### 3.1 Urine proteome analysis

#### 3.1.1 Identification of urinary proteins

14 samples from the experimental and control groups collected after intragastric administration were analyzed by LC-MS/MS. A total of 1542 proteins were identified according to the criteria that each protein contains at least 2 specific peptides and FDR<1% at the protein level. Urinary proteins of the two groups were compared. 166 differential proteins were identified under the screening conditions of FC ≥ 1.5 or ≤ 0.67 and *P* < 0.05, of which 88 were down-regulated and 78 were up-regulated. Detailed information on the differential proteins was listed in Supplementary Table 1.

Hierarchical cluster analysis was performed on the identified total proteins and differential proteins respectively (Figure 2), and principal component analysis was performed on the identified differential proteins (Figure 3), which could distinguish the samples of the experimental and control groups.

**Figure 2.**
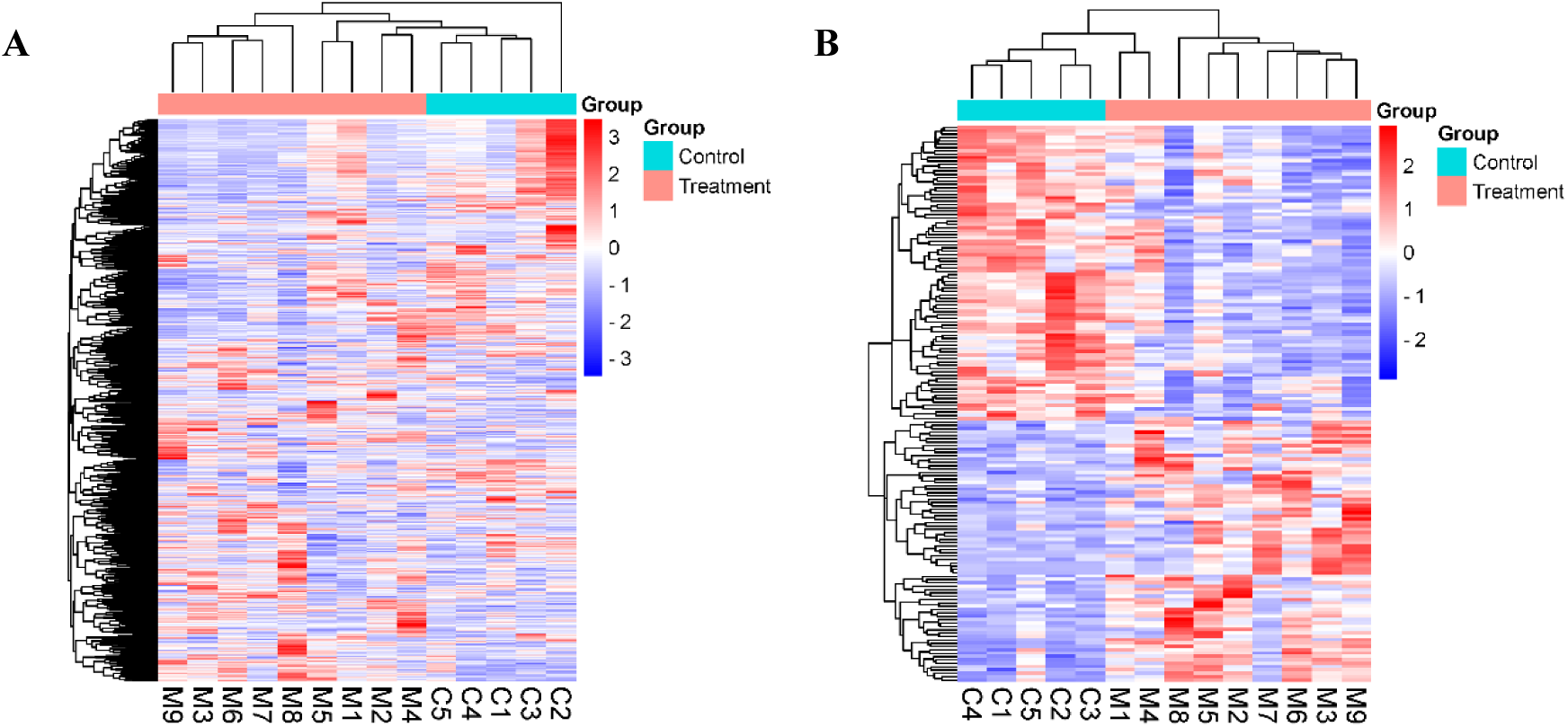
Hierarchical cluster analysis of total and differential proteins. (A) total proteins; (B) differential proteins.

**Figure 3.**
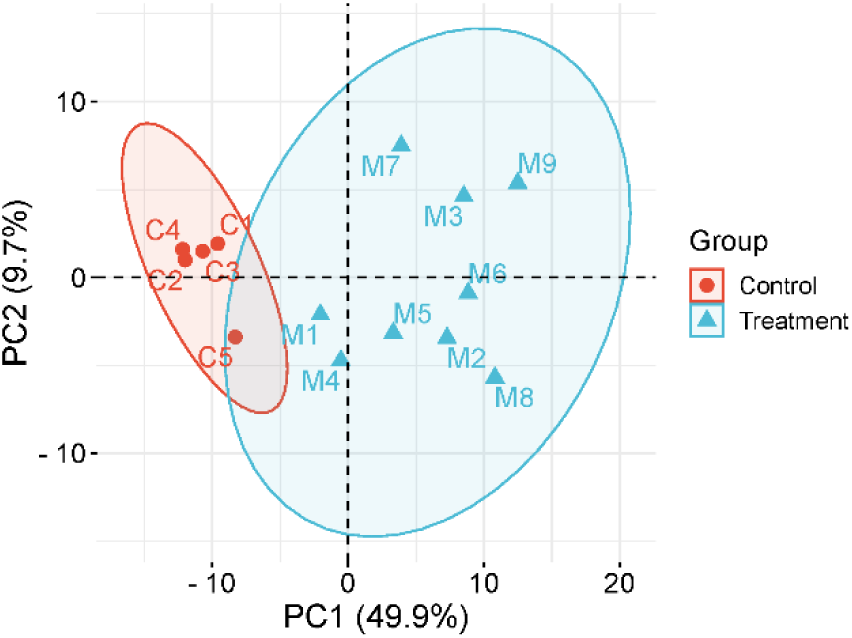
Principal component analysis of differential proteins.

#### 3.1.2 Randomized grouping test for total proteins

To determine the possibility of random generation of identified differential proteins, the total proteins identified from 14 samples in two groups were verified by a randomized grouping test. 14 samples were disrupted and randomly combined into two new groups, with a total of 2002 combination cases, which were screened for differences according to the same criteria (FC ≥ 1.5 or ≤ 0.67, *P* < 0.05). The average number of differential proteins yielded was 38.46, indicating that at least 76.83% of the differential proteins were not randomly generated. The results showed that the 166 different proteins identified were reliable.

#### 3.1.3 Analysis of differential proteins

Functional analysis of 166 differential proteins was performed using the PubMed database. 4 proteins have been reported to be directly affected by metformin. Other members of two protein families have been reported to be directly affected by metformin. 27 proteins, although have not been reported to be directly affected by metformin, are functionally related to metformin efficacy. Detailed information on each protein is as follows:

##### 3.1.3.1 Differential proteins reported to be directly affected by metformin

###### (1) Pyruvate kinase

Metformin enhances pyruvate kinase activity in hepatocytes and inhibits the process of gluconeogenesis [30]. Inhibition of hepatic gluconeogenesis is one of the main ways that metformin exerts a hypoglycemic effect [31]. Some researchers have proposed the hypothesis that metformin potentiates the allosteric activation of pyruvate kinase by fructose-1,6-diphosphate, suggesting pyruvate kinase as the locus of metformin’s clinical action [8].

**Table 1.** Differential proteins reported to be directly affected by metformin.

| UniProt ID | Protein name | Trend | FC | P-value | Ref. |
| --- | --- | --- | --- | --- | --- |
| A0A0H2UI07 | Pyruvate kinase | ↓ | 0.54 | 3.44E-05 | [8,30-36] |
| G3V8K5 | Growth differentiation factor 15 | ↑ | 2.01 | 3.70E-04 | [37-42] |
| A0A0G2K3C5 | Cystathionine gamma-lyase | ↓ | 0.50 | 1.06E-03 | [43,44] |
| F1LMN1 | Cytochrome P450 | ↑ | 13.22 | 1.52E-02 | [45] |

In addition, pyruvate kinase M2 (PKM2), a downstream molecule in the PI3K/AKT/mTOR signaling pathway, is overexpressed in nearly all tumor cells and plays a crucial role in the Warburg effect [32,33]. Metformin downregulates PKM2 expression in gastric cancer cells [34], esophageal cancer cells [35], and breast cancer cells [36]. Modulating the activity or expression of PKM2 has been reported to be a promising strategy to enhance the anticancer effect of metformin.

###### (2) Growth differentiation factor 15 (GDF15)

Several studies have shown that taking metformin increases GDF15 levels [37–40]. GDF15 is a cytokine with anti-inflammatory properties that increases insulin sensitivity, suppresses appetite, reduces body weight in both diabetic and non-diabetic patients, and improves the prognosis of diabetic patients. GDF15 levels have also been associated with the progression of diabetic complications, including thrombosis, diabetic nephropathy, diabetic neuropathy degeneration, and diabetic retinopathy [41]. GDF15 levels have been reported to be a novel biomarker of metformin administration in patients with abnormal blood glucose, and its concentration may reflect the dose of metformin [38]. In addition, due to its anti-inflammatory and appetite suppressant effects, GDF15 has great potential in the treatment of a variety of metabolic disorders such as obesity, T2DM, non-alcoholic fatty liver disease, cardiovascular disease, and cancer cachexia [42].

###### (3) Cystathionine gamma-lyase (CSE)

Metformin can alleviate atherosclerosis by regulating CSE expression and promoting hydrogen sulfide (H₂S) production [43]. In rats exposed to bisphenol A (BPA), metformin upregulates the expression of CSE and cystathionine β-synthase (CBS), reduces serum homocysteine levels, and protects against BPA-induced hepatic injury [44].

###### (4) Cytochrome P450

In rats, metformin is metabolized mainly by CYP2C11, 2D1, and 3A1/2, which are hepatic microsomal cytochrome P450 (CYP) isozymes [45].

##### 3.1.3.2 Other members of the protein family have been reported to be directly affected by metformin

###### (1) Solute carrier family 22, member 21

Solute carrier family 22 is an organic cation transporter protein involved in transporting various endogenous and exogenous substances. The main transporter proteins of metformin are solute carrier family 22 member 1 (OCT1) and member 4 (OCTN1) [46]. Among them, genetic polymorphism of OCT1 can affect the pharmacokinetics of metformin and gastrointestinal intolerance, thus affecting the individual response to metformin [47–49].

**Table 2.** Other members of the family of differential proteins reported to be directly affected by metformin.

| UniProt ID | Protein name | Trend | FC | P-value | Ref. |
| --- | --- | --- | --- | --- | --- |
| A0A0G2JVF2 | Solute carrier family 22 | ↓ | 0.24 | 4.66E-02 | [46-49] |
| D3ZHS5 | Carboxypeptidase A4 | ↑ | 49.79 | 8.39E-03 | [50] |

###### (2) Carboxypeptidase A4

Carboxypeptidase A4 (FC = 49.79, *p* = 8.39×10^−3^) had the largest intergroup FC value among the 166 differential proteins identified in this study. Genetic variation in carboxypeptidase A6 (CPA6) has been reported to be associated with metformin response in patients with T2DM [50].

##### 3.1.3.3 Not reported to be directly affected by metformin, but protein function is associated with metformin efficacy

###### (1) Secreted Ly6/Plaur domain containing 2(SLURP-2)

SLURP-2 (FC = 1.54, *p* = 5.67×10^−4^) ranked as the 5^th^ smallest *p*-value among all differential proteins identified in this study. SLURP-2 is a new member of the Ly-6 superfamily with up-regulated expression in psoriasis patients and may be involved in the pathophysiological process of psoriasis through keratinocyte proliferation and T cell differentiation/activation [51]. Several studies have shown that metformin is effective in improving treatment outcomes and metabolic syndrome in patients with psoriasis [52–54], and that long-term use of metformin is also associated with a reduced risk of psoriasis [55].

**Table 3.** Differential proteins not reported to be directly affected by metformin but functionally associated with metformin efficacy.

| UniProt ID | Protein name | Trend | FC | P-value | Effect/Associated with a disease | Ref. |
| --- | --- | --- | --- | --- | --- | --- |
| D3ZUR5 | Secreted Ly6/Plaur domain containing 2 | ↑ | 1.54 | 5.67E-04 | psoriasis | [51-55] |
| A0A0G2J SI5 | Chymotrypsin-like elastase family member 1 | ↑ | 25.11 | 3.91E-02 | emphysema | [56,57] |
| Q63471 | BPI fold-containing family A member 2 | ↑ | 24.76 | 4.66E-03 | acute kidney injury | [58,59] |
| Q5XIM9 | T-complex protein 1 subunit beta (TCP-1-beta) | ↓ | 0.36 | 6.86E-04 | diabetic nephropathy | [60-63] |
| G3V709 | Nicotinate phosphoribosyltransferase | ↓ | 0.42 | 4.67E-03 | aging | [64] |
| A0A0H2 UI19 | Coagulation factor XII | ↑ | 1.51 | 2.91E-03 | coagulation factor | [65] |
| A0A0H2 UHR6 | Coagulation factor X | ↑ | 2.29 | 2.08E-04 | coagulation factor | [65] |
| Q07009 | Calpain-2 catalytic subunit | ↓ | 0.21 | 9.08E-04 | atrial fibrillation | [68] |
| A0A0G2J<br>VZ6 | Integrin subunit alpha V | ↓ | 0.27 | 4.92E-02 | high blood<br>glucose/diabetes | [69] |
| P97608 | 5-oxoprolinase | ↓ | 0.27 | 2.41E-02 | high blood<br>glucose/diabetes | [70] |
| Q4KLZ6 | Triokinase/FMN cyclase | ↓ | 0.61 | 2.45E-02 | high blood<br>glucose/diabetes | [71] |
| A0A0H2<br>UHE4 | Regenerating family<br>member 3 beta | ↑ | 4.26 | 2.99E-02 | high blood<br>glucose/diabetes | [72] |
| D4A0W2 | Lysozyme f1 | ↑ | 19.23 | 3.68E-03 | high blood<br>glucose/diabetes | [73] |
| P07824 | Arginase-1 | ↑ | 19.52 | 1.61E-02 | high blood<br>glucose/diabetes | [74] |
| P19629 | L-lactate dehydrogenase C<br>chain (LDH-C) | ↓ | 0.08 | 2.37E-02 | cancer | [81,82] |
| Q6TA48 | Mucosal pentraxin | ↓ | 0.12 | 4.38E-02 | cancer | [83-87] |
| F1LRT9 | Dynein cytoplasmic 1<br>heavy chain 1 | ↓ | 0.21 | 1.59E-03 | cancer | [88-90] |
| P16446 | Phosphatidylinositol<br>transfer protein alpha<br>isoform | ↓ | 0.23 | 3.48E-02 | cancer, Duchenne<br>muscular<br>dystrophy | [91-93] |
| D3Z9E5 | Sodium-coupled<br>monocarboxylate<br>transporter 1 | ↓ | 0.36 | 5.73E-04 | cancer | [94-97] |
| F1M3L7 | Epidermal growth factor<br>receptor kinase substrate 8 | ↓ | 0.58 | 5.68E-04 | cancer | [98-103] |
| Q66H12 | Alpha-N-acetylgalactosaminidase | ↓ | 0.65 | 6.84E-04 | cancer | [104] |
| Q4FZU6 | Annexin A8 | ↑ | 4.07 | 2.72E-02 | cancer | [105,106] |
| P02793 | Ferritin light chain 1 | ↑ | 4.48 | 1.97E-02 | cancer | [76,107] |
| F1LQQ8 | Beta-glucuronidase | ↑ | 8.86 | 6.67E-03 | cancer | [108] |
| Q6AYR8 | Secernin-2 | ↓ | 0.25 | 1.03E-02 | cognitive<br>dysfunction | [116] |
| Q3KR94 | Vitronectin | ↑ | 1.53 | 5.99E-04 | cognitive<br>dysfunction | [117] |
| F1M7F7 | Complement component<br>C6 | ↑ | 1.96 | 2.66E-05 | cognitive<br>dysfunction | [118,119] |

###### (2) Chymotrypsin-like elastase family member 1

Chymotrypsin-like elastase family member 1 (FC = 25.11, *p* = 3.91×10^−2^) has the second highest FC value of all differential proteins identified in this study after carboxypeptidase A4. The gene encoding this protein, *Cela1*, is expressed during lung development and is closely associated with the process of stretch-dependent remodeling of the lungs physiologically and pathologically. Its expression is increased in both mice and humans with α-1 antitrypsin-deficient emphysema, which may be a specific target for treating α-1 antitrypsin-deficient emphysema [56]. Metformin has a potentially important role in the treatment of emphysema in mice and humans, especially in slowing down the progression of emphysema [57].

###### (3) BPI fold-containing family A member 2(BPIFA2)

BPIFA2 levels are higher in the blood and urine of patients with acute kidney injury compared to healthy individuals, and BPIFA2 can be used as an early biomarker of acute kidney injury [58]. Even the administration of low doses of metformin exacerbates renal ischemia-reperfusion-induced acute kidney injury and increases mortality in mice [59].

###### (4) T-complex protein 1 subunit beta (TCP-1β)

TCP-1β can be used as a biomarker for the glomerular hyperfiltration phase of type 2 diabetic nephropathy [60]. Several studies have shown that metformin plays an important role in alleviating the occurrence of diabetic nephropathy [61–63].

###### (5) Nicotinate phosphoribosyltransferase

Nicotinate phosphoribosyltransferase plays an important role in the metabolism of trigonelline, which is used in the synthesis of nicotinamide adenine dinucleotide (NAD+), enhances mitochondrial activity, and has great potential for amelioration of age-related muscle decline [64].

###### (6) Coagulation factor X, coagulation factor XII

Diabetic patients exhibit a hypercoagulable state compared to healthy individuals, which is associated with high levels of coagulation factors (II, V, VII, VIII, and X) and low levels of anticoagulants (protein C) [65].

###### (7) Calpain-2 catalytic subunit

Several studies have shown that metformin plays a positive role in cardiovascular protection. Metformin reduces cardiovascular disease mortality, all-cause mortality, and the risk of cardiovascular events in patients with coronary artery disease [66]. In patients with T2DM, treatment with metformin is associated with a reduction in cardiovascular mortality and morbidity [4]. Metformin also protects the heart against hypertrophic and apoptotic remodeling after myocardial infarction [67]. Calpain 2 has been reported to be significantly elevated in atrial samples from patients with atrial fibrillation compared to those in sinus rhythm, and calpain 2 may be associated with the development of atrial fibrillation in patients with heart valve disease and diabetes [68].

Proteins associated with high blood glucose or diabetes are as follows:

###### (1) Integrin subunit alpha V

Hyperglycemia reduces integrin subunits alpha v and alpha 5 on the surface of dermal fibroblasts, which affects fibroblast migration and the wound healing process. This may be one of the mechanisms of defective wound healing in diabetes [69].

###### (2) 5-oxoprolinase (OPLAH)

Down-regulation of OPLAH in human skeletal muscle cells (HSKMCs) may lead to insulin resistance and glucose uptake impairment through oxidative stress, which may become a new therapeutic target for T2DM [70].

###### (3) Triokinase/FMN cyclase

As a new NRF2 target gene, *Tkfc* encoding triokinase/FMN cyclase is up-regulated in atypical NRF2 activation and plays an important role in promoting hepatic fructose metabolism and gluconeogenesis, and influencing blood glucose homeostasis [71].

###### (4) Regenerating family member 3 beta (Reg3β)

Recombinant Reg3β protein is important in preventing streptozotocin-induced diabetes and pancreatic islet beta cell damage in mice [72].

###### (5) Lysozyme f1

In patients with T2DM, the structure and function of lysozyme are altered. L-lysine, as a chemical molecular chaperone, significantly improves lysozyme structure and function, reverses glycosylation changes, increases lysozyme activity, and helps prevent diabetic complications [73].

###### (6) Arginase-1 (ARG1)

Arginase 1 (ARG1) and Arginase 2 (ARG2) play important roles in regulating β-cell function, insulin resistance (IR), and vascular complications by regulating L-arginine metabolism, nitric oxide (NO) production, inflammatory responses, and oxidative stress. Abnormal alterations in arginase expression and activity are closely associated with the development of diabetes and its complications. Targeting arginase is expected to be a therapeutic approach for diabetes [74].

In addition, cancer has become the leading cause of death in diabetic patients in high-income countries [75]. A meta-analysis study involving 10,695,875 patients with T2DM showed that patients using metformin had a significantly lower risk of cancer compared to other glucose-lowering medications. Subgroup analyses showed a significant reduction in the risk of bladder cancer, colorectal cancer, gastric cancer, liver cancer, lung cancer, pancreatic cancer, and prostate cancer with the use of metformin [3]. Several other studies have shown that metformin has great potential in the treatment of glioma [76], cervical cancer [77], acute myeloid leukemia [78], breast cancer [79], and ovarian cancer [80]. The differential proteins associated with cancer are as follows:

###### (1) L-lactate dehydrogenase C chain, LDH-C

L-lactate is generated by the reduction of pyruvate catalyzed by lactate dehydrogenase. The level of lactate dehydrogenase 5 (LDH5) is higher than that of normal tissues in many human tumor tissues. *LDHC* is also expressed in a variety of tumors, including lung cancer, melanoma, prostate cancer, and breast cancer [81,82]. The enzymes involved in L-lactate metabolism are closely related to the pathophysiological processes of diabetes, cancer, and other diseases, which may provide new strategies and approaches for disease treatment [81].

###### (2) Mucosal pentraxin

Heme, which has been reported to be associated with red meat and the risks of colon cancer, can lead to more than a 10-fold downregulation of *Mptx* in rats [83]. *Mptx* may be involved in processing damaged cells in the colonic mucosa. Its expression is associated with colonic cell renewal, which may be a marker for diet-induced stress in the colonic mucosa [83,84]. Meta-analysis studies have shown that metformin use is associated with a reduced risk of developing colon cancer [85], with a significant reduction in overall colon cancer mortality, and with a better prognosis for colon cancer patients [86,87].

###### (3) Dynein cytoplasmic 1 heavy chain 1 (DYNC1H1)

DYNC1H1 encodes the cytoplasmic dynein heavy chain family, which links phagocytosis to apoptosis and prevents various diseases, including cancer, neurodegenerative diseases, and autoimmune diseases [88,89]. Studies have shown that DYNC1H1 is a new prognostically relevant biomarker for hepatocellular carcinoma, which is associated with epithelial-mesenchymal transition and immune infiltration, and has great potential for early diagnosis and effective intervention in hepatocellular carcinoma [88]. Metformin may influence the early progression of hepatocellular carcinoma associated with non-alcoholic fatty liver disease/non-alcoholic steatohepatitis (NAFLD/NASH) by modulating macrophage polarization and T-cell infiltration [90].

###### (4) Phosphatidylinositol transfer protein alpha isoform (PITP-alpha)

PITP is an abundant and ubiquitous soluble protein. Increased expression of PITPα/β in gastric cancer tissues is associated with poor prognosis of gastric cancer, making it a potential target for cancer treatment [91].

In addition, decreased expression of PITPα is related to the pathological improvement of Duchenne muscular dystrophy and is a potential target for the treatment of Duchenne muscular dystrophy [92]. Metformin improves muscle function and reduces neuromuscular deficits in mice with muscular dystrophy, and has the potential to be used as a therapeutic agent for patients with Duchenne muscular dystrophy [93].

###### (5) Sodium-coupled monocarboxylate transporter 1

In various cancers, *SLC5A8*, the gene encoding sodium-coupled monocarboxylate transporter 1, can function as a tumor suppressor gene. In cervical cancer, *SLC5A8* is silenced by DNA hypermethylation and histone deacetylation, and can serve as a biomarker and therapeutic target for the diagnosis and prognosis of cervical cancer [94]. *SLC5A8* has been identified as an oncogene associated with colorectal cancer and is methylated and silenced in colon cancer [95]. Among hypermethylated and silencing genes associated with acute myeloid leukemia in mixed lineage leukemia partial tandem duplication (*MLL*-PTD), silencing of the tumor suppressor gene *SLC5A8* promotes leukemia occurrence [96]. *SLC5A8* is silenced in mouse breast cancer, and reactivation of *SLC5A8* expression may be a new therapeutic strategy for breast cancer [97].

###### (6) Epidermal growth factor receptor kinase substrate 8 (Eps8)

Eps8 is highly expressed in various human tumor types, including colorectal cancer [98], pituitary tumors [99], oral squamous cell carcinoma [100], esophageal cancer [101], pancreatic cancer [102], and cervical cancer [103], and participates in many signaling pathways associated with carcinogenesis, metastasis, and proliferation. Eps8 is a biomarker of poor prognosis in cancer patients.

###### (7) Alpha-N-acetylgalactosaminidase

Down-regulation of α-N-acetylgalactosaminidase expression mediated by shRNA inhibits migration and invasion of breast and ovarian cancer cell lines. α-N-acetylgalactosaminidase is expected to be an anti-cancer therapeutic target [104].

###### (8) Annexin A8

Studies have shown that annexin A8 is overexpressed in pancreatic cancer [105]. Its increased expression is associated with poor prognosis in early-stage pancreatic cancer and can be used as a prognostic marker and potential therapeutic target in pancreatic cancer [106].

###### (9) Ferritin light chain 1, FTL

Hypoxia-induced FTL is a regulator of epithelial mesenchymal transition, which can be used as a prognostic marker for glioma and a novel biomarker for the response of anti-tumor drug temozolomide [107]. Metformin inhibits glioma cell stemness and epithelial-mesenchymal transition by modulating the activity of YAP, a key effector of the Hippo pathway [76].

###### (10) Beta-glucuronidase

High levels of urinary beta-glucuronidase are observed in patients with bladder cancer [108].

Cognitive dysfunction has also been reported as one of the many complications of diabetes mellitus [109]. Several studies have shown that diabetic patients are at an increased risk of developing dementia such as Alzheimer’s disease [110–113]. Metformin plays a positive role in ameliorating cognitive impairment and alleviating memory loss [2,12,114,115]. The differential proteins associated with cognitive dysfunction are as follows:

###### (1) Secernin-2

*SCRN2*, *LCMT1*, *LRRC46*, *MRPL10*, *SP6*, *OSBPL7* are significantly associated with Aβ standardized uptake value ratio in the brain. single nucleotide polymorphisms (SNPs) on these genes are also associated with reduced hippocampal volume and cognitive scores. These six genes may be new therapeutic targets for Alzheimer’s disease [116].

###### (2) Vitronectin (VTN)

VTN is a multifunctional glycoprotein. VTN and its receptors have been associated with various diseases, including tumors, coagulation disorders, inflammatory diseases, and a variety of neurodegenerative disorders. VTN plays an important role in neuronal function and neurodegenerative diseases, and can be involved in neural differentiation, neuro-nutrition, and neurogenesis, regulating axon size, supporting and guiding neuronal extension. Additionally, VTN protects the brain by interacting with integrin receptors in vascular endothelial cells to reduce the permeability of the blood-brain barrier [117].

###### (3) Complement component C6

Complement component C6 (FC = 1.96, *p* = 2.66×10^−5^) has the smallest *p*-value among all differential proteins identified in this study. Complement is an important factor in the progression of neurodegenerative diseases such as Alzheimer’s disease, amyotrophic lateral sclerosis, and schizophrenia [118]. Activation of the innate immune response, especially the terminal pathway of the complement system, can lead to the formation of the membrane attack complex (MAC) and delay peripheral nervous system regeneration, which is a key cause of neuronal damage. Complement component C6 is important in the activation of the complement system and MAC formation. And C6 deficiency facilitates neuronal recovery after trauma [119].

#### 3.1.4 Biological pathway analysis

Biological process and molecular function enrichment analysis of the identified differential proteins were performed using the DAVID database (Figure 4).

**Figure 4.**
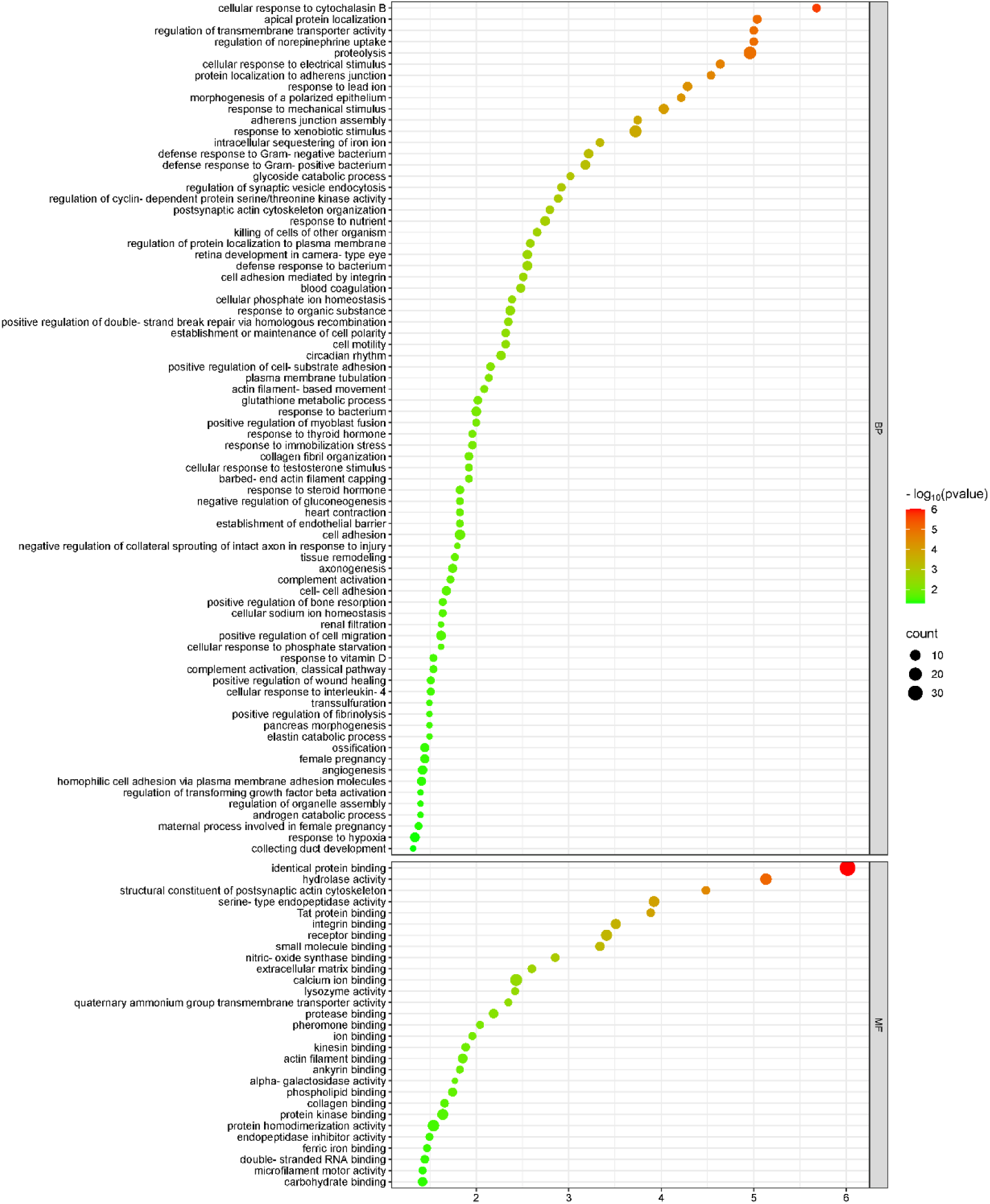
Enrichment analysis of biological processes and molecular functions of identified differential proteins.

These differential proteins were mainly involved in biological processes including cellular response to cytochalasin B, apical protein localization, regulation of norepinephrine uptake, regulation of transmembrane transporter protein activity, proteolysis, glycoside catabolic process, regulation of cyclin-dependent protein serine/threonine kinase activity, blood coagulation, glutathione metabolic process, establishment of endothelial barrier, and negative regulation of gluconeogenesis. Among them, blood coagulation, glutathione metabolic process, establishment of endothelial barrier, and negative regulation of gluconeogenesis have been reported to be associated with metformin action. Diabetic patients exhibit hypercoagulability compared to healthy individuals [65]. Metformin modulates glutathione metabolism and influences the progression of thyroid cancer [120]. Vascular endothelium is closely related to the regulation of cardiovascular function. Several studies have shown that metformin is important in improving endothelial function [121–124]. Inhibition of gluconeogenesis is one of the main pathways through which metformin exerts its hypoglycemic effect [31].

Among the molecular functions, most of these differential proteins were found to have functions such as identical protein binding, hydrolase activity, structural constituent of the postsynaptic actin cytoskeleton, serine-type endopeptidase activity, Tat protein binding, integrin binding, receptor binding, small molecule binding, nitric-oxide synthase binding, and lysozyme activity. Studies have shown that nitric-oxide synthase plays an important role in oxidative stress and vascular diseases [125]. The uncoupling of endothelial nitric oxide synthase (eNOS) may explain the pathogenesis of diabetic vascular diseases caused by decreased levels and impaired function of endothelial progenitor cells to a certain extent [126]. In addition, metformin promotes vascular regeneration and neurological recovery after spinal cord injury in aged mice by activating the AMPK/eNOS signaling pathway [127]. In patients with T2DM, the structure and function of lysozyme are altered. Reversing the changes caused by glycosylation and increasing lysozyme activity contribute to the prevention of diabetic complications [73].

Kyoto Encyclopedia of Genes and Genomes (KEGG) enrichment analysis showed that significant enrichment pathways include complement and coagulation cascades, proteoglycans in cancer, focal adhesion, regulation of actin cytoskeleton, dilated cardiomyopathy, arrhythmogenic right ventricular cardiomyopathy, ECM-receptor interaction, hypertrophic cardiomyopathy, glycerolipid metabolism, platelet activation, and PI3K-Akt signaling pathway (Figure 5). Among them, glycerolipid metabolism, platelet activation, and PI3K-Akt signaling pathways have been reported to be associated with metformin efficacy. T2DM induces inactivation of the glycerolipid metabolism pathway, and vanillin with antidiabetic activity significantly reverses this change [128]. Platelet surface receptors and platelet activation markers in T2DM patients significantly differed from those in healthy individuals [129]. Platelet activation has also been associated with the development of chronic diseases such as atherosclerosis, coronary artery disease, and cerebrovascular disease [130]. The PI3K-Akt signaling pathway is the most frequently activated in cancer. Under physiological conditions, this pathway is activated under the action of insulin, growth factors, and cytokines, regulates key metabolic processes such as glucose metabolism and macromolecular biosynthesis, and maintains systemic metabolic homeostasis [131].

**Figure 5.**
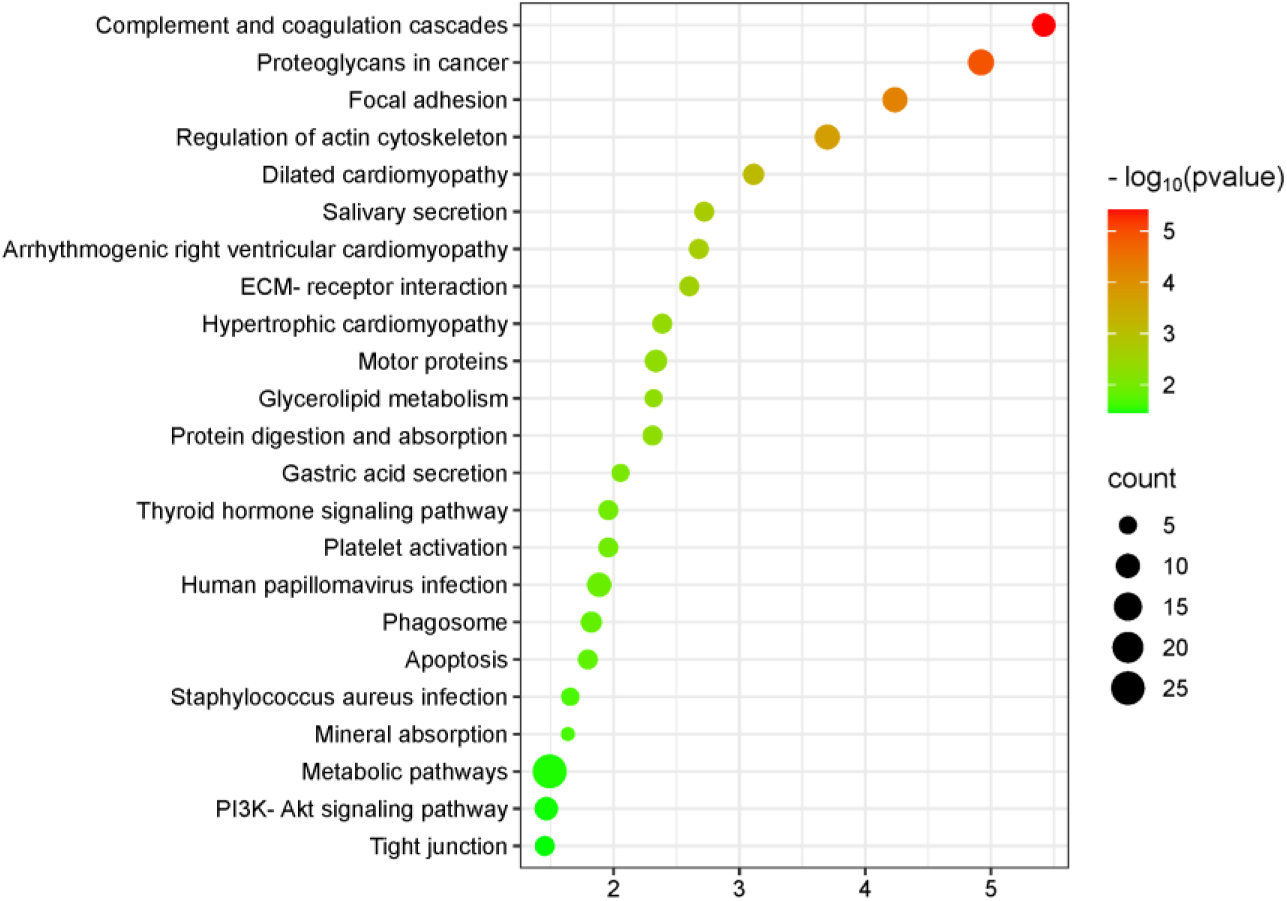
Enrichment analysis of the KEGG pathways of identified differential proteins.

### 3.2 Analysis of urine proteome modifications

#### 3.2.1 Identification of differentially modified peptides

Relying on the non-labeled quantitative proteome method, the data from 14 samples were obtained by LC-MS/MS analysis. A search based on open-pFind yielded detailed information on the mass spectra number of modified peptides in each sample, including the proteins in which the peptides are located and the types of modifications presented in the peptides. The modified peptides, which had a reproducibility of over 50%, were screened respectively in the experimental and control groups, and then the union set was taken. A total of 3206 modified peptides were identified. 285 differentially modified peptides were identified under the screening conditions of FC ≥ 1.5 or ≤ 0.67 and *P* < 0.05. Detailed information was listed in Appendix 1, including the peptide sequences, modification types, and proteins in which the differentially modified peptides are located.

Hierarchical cluster analysis was performed on the identified total modified peptides and differentially modified peptides respectively (Figure 6), and principal component analysis was performed on the differentially modified peptides (Figure 7), which could distinguish the samples of the experimental and control groups. The results of principal component analysis (Figures 3 and 7) for both the differential proteins and the differentially modified peptides showed that the sample points in the experimental group were highly dispersed compared to the control group, indicating some inter-individual variation. It has been reported that SNPs in the genes encoding metformin transporters such as OCT1, OCT2, MATE1, and MATE2 are significantly associated with the efficacy and toxicity of metformin [132]. In addition, individual response to metformin treatment is also influenced by factors including DNA methylation level [133], individual health level [134,135], and gender [136]. This suggests the need to pay attention to the effects of individual differences in drug research.

**Figure 6.**
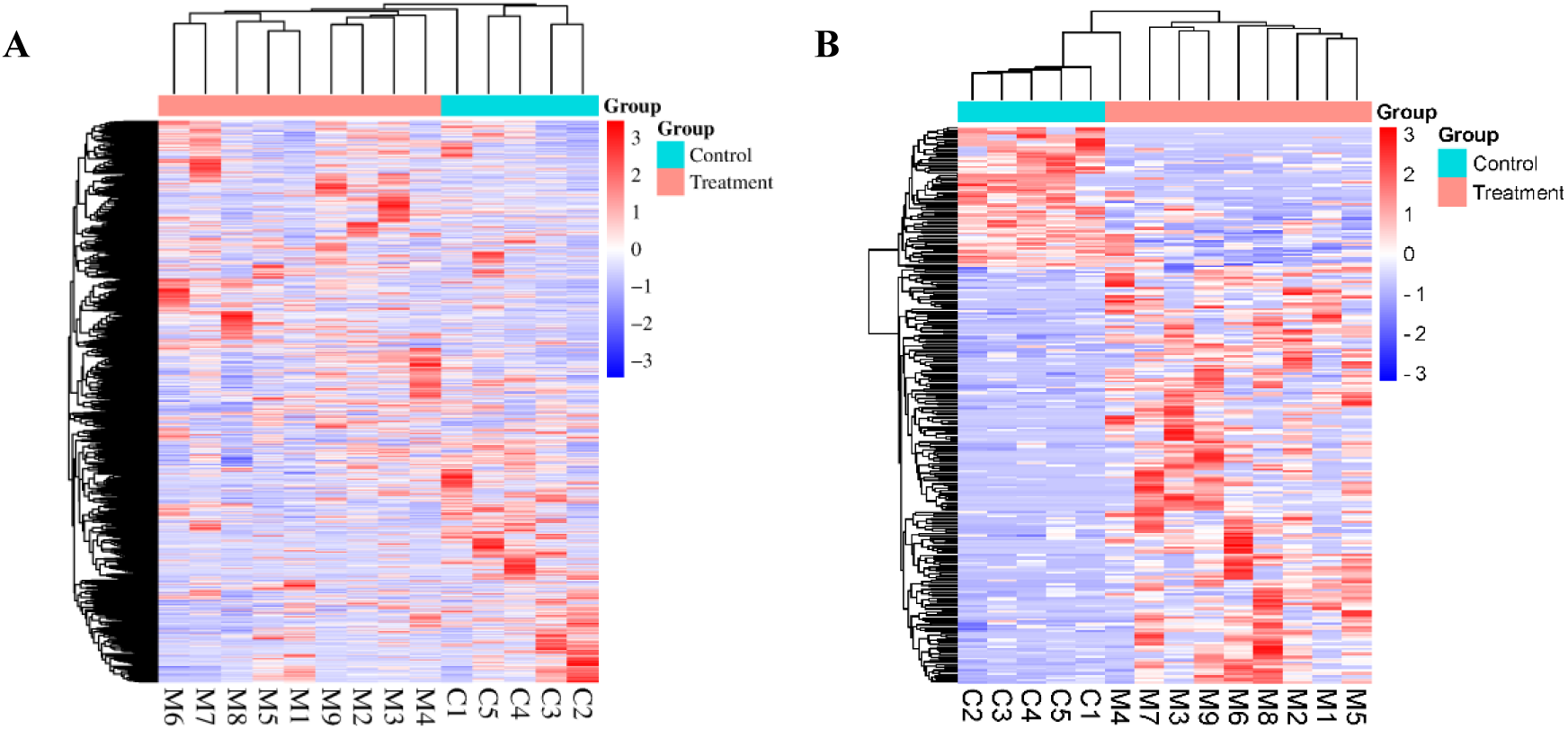
Hierarchical cluster analysis of total and differentially modified peptides. (A) total modified peptides; (B) differentially modified peptides.

**Figure 7.**
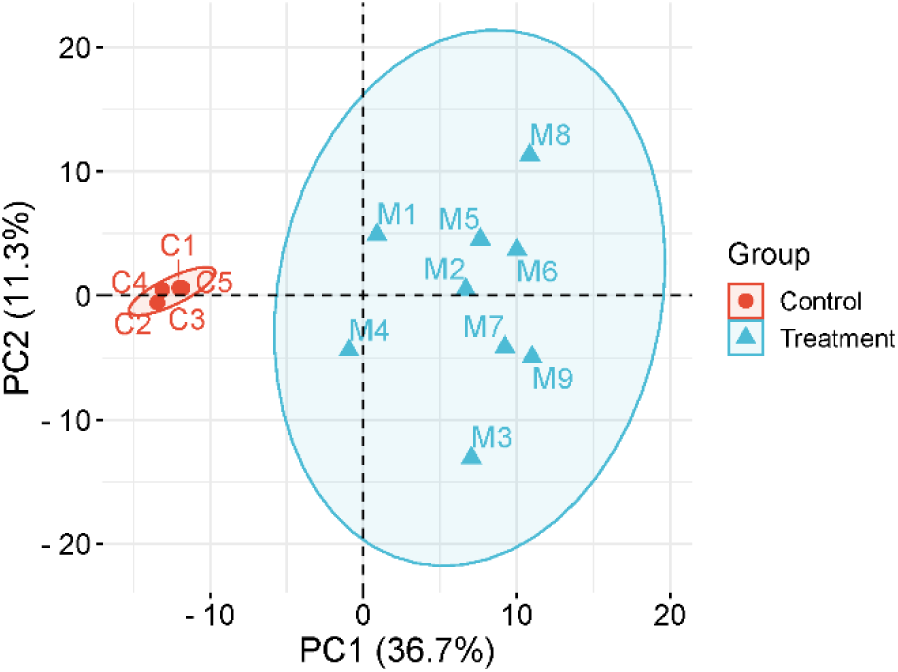
Principal component analysis of differentially modified peptides.

#### 3.2.2 Randomized grouping test for total modified peptides

To determine the possibility of random generation of identified differentially modified peptides, the total modified peptides were verified by a randomized grouping test. 14 samples were disrupted and randomly combined into two new groups, with a total of 2002 combination cases, which were screened for differences according to the same criteria (FC ≥ 1.5 or ≤ 0.67, *P* < 0.05). The average number of differentially modified peptides yielded was 132.8, indicating that at least 53.4% of the differentially modified peptides were not randomly generated.

#### 3.2.3 Analysis of biological pathways enriched in proteins where differentially modified peptides are located

A total of 127 proteins containing differentially modified peptides were screened, details of which were shown in Appendix 2. Biological process and molecular function enrichment analysis of the proteins containing differentially modified peptides were performed using the DAVID database (Figure 8).

**Figure 8.**
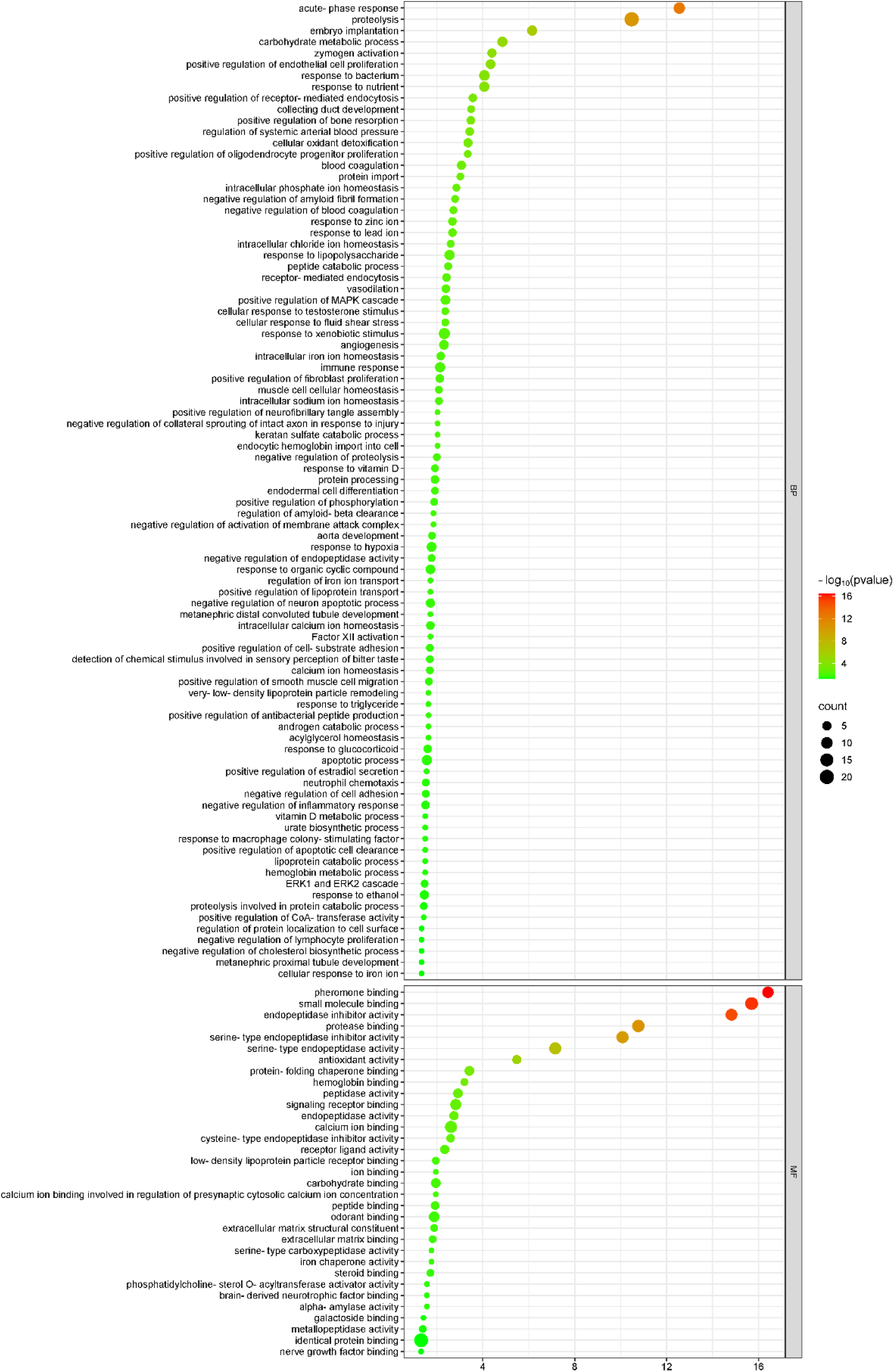
Enrichment analysis of biological processes and molecular functions of proteins containing differentially modified peptides.

Proteins containing differentially modified peptides were mainly involved in biological processes including acute-phase response, proteolysis, carbohydrate metabolic process, zymogen activation, positive regulation of endothelial cell proliferation, response to bacterium, response to nutrient, positive regulation of receptor-mediated endocytosis, regulation of systemic arterial blood pressure, cellular oxidant detoxification, positive regulation of oligodendrocyte progenitor proliferation, blood coagulation, intracellular phosphate ion homeostasis, negative regulation of amyloid fibril formation, and negative regulation of blood coagulation.

Among the molecular functions, most of these proteins were found to have functions such as pheromone binding, small molecule binding, endopeptidase inhibitor activity, protease binding, serine-type endopeptidase inhibitor activity, serine-type endopeptidase activity, antioxidant activity, protein-folding chaperone binding, and hemoglobin binding.

KEGG enrichment analysis showed that significantly enriched pathways include complement and coagulation cascades, lysosome, renin-angiotensin system, staphylococcus aureus infection, and protein digestion and absorption (Figure 9). Angiotensin II, a key component of the renin-angiotensin system (RAS), is an important target for effectively lowering blood pressure and preventing cardiovascular disease and kidney injury progression in diabetic patients [137].

**Figure 9.**
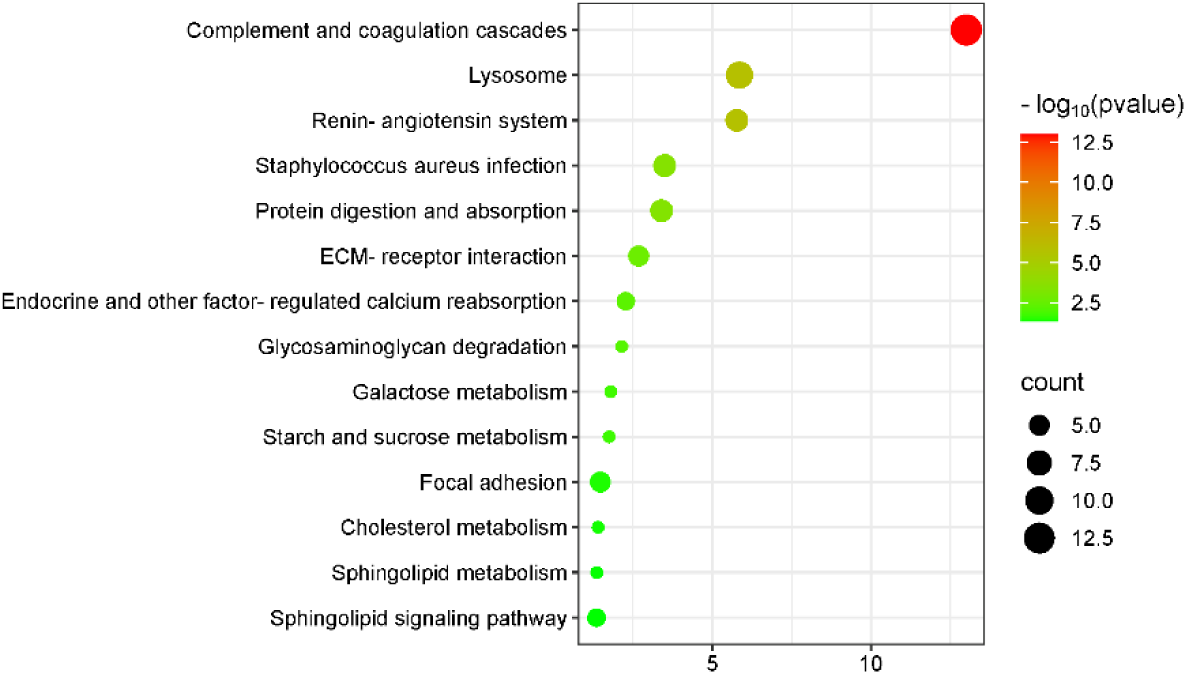
Enrichment analysis of the KEGG pathway of proteins containing differentially modified peptides.

#### 3.2.4 Differentially modified peptides from presence to absence or absence to presence

The differentially modified peptides with significant changes were screened with the following criteria: identified in over half of the control group samples but none in the experimental group, or vice versa. 100 such peptides were screened, 9 identified in over half of the samples in the control group but not in the experimental group, and 91 identified in over half of the samples in the experimental group but not in the control group. Detailed information was listed in Appendix 3, including the peptide sequences, modification types, the proteins where they are located, and the number of the modified peptide mass spectra of each sample.

A total of 57 proteins containing differentially modified peptides from presence to absence or absence to presence were screened, details of which were shown in Appendix 4. Biological process and molecular function enrichment analysis of these proteins were performed using the DAVID database (Figure 10). These proteins were mainly involved in biological processes including proteolysis, zymogen activation, response to lipopolysaccharide, cellular oxidant detoxification, regulation of systemic arterial blood pressure, acute-phase response, response to nutrient, negative regulation of activation of membrane attack complex, positive regulation of lipoprotein transport, acylglycerol homeostasis, very-low-density lipoprotein particle remodeling, response to triglyceride, lipoprotein catabolic process, positive regulation of CoA-transferase activity, reverse cholesterol transport, peripheral nervous system axon regeneration, negative regulation of lipid biosynthetic process, and high-density lipoprotein particle remodeling. Long-term use of metformin has been reported to reduce cholesterol and low-density lipoprotein levels in mice [138]. Among the molecular functions, most of these proteins have functions such as pheromone binding, small molecule binding, endopeptidase inhibitor activity, antioxidant activity, serine-type endopeptidase activity, serine-type endopeptidase inhibitor activity, phosphatidylcholine-sterol O-acyltransferase activator activity, hemoglobin binding, and carbohydrate binding.

**Figure 10.**
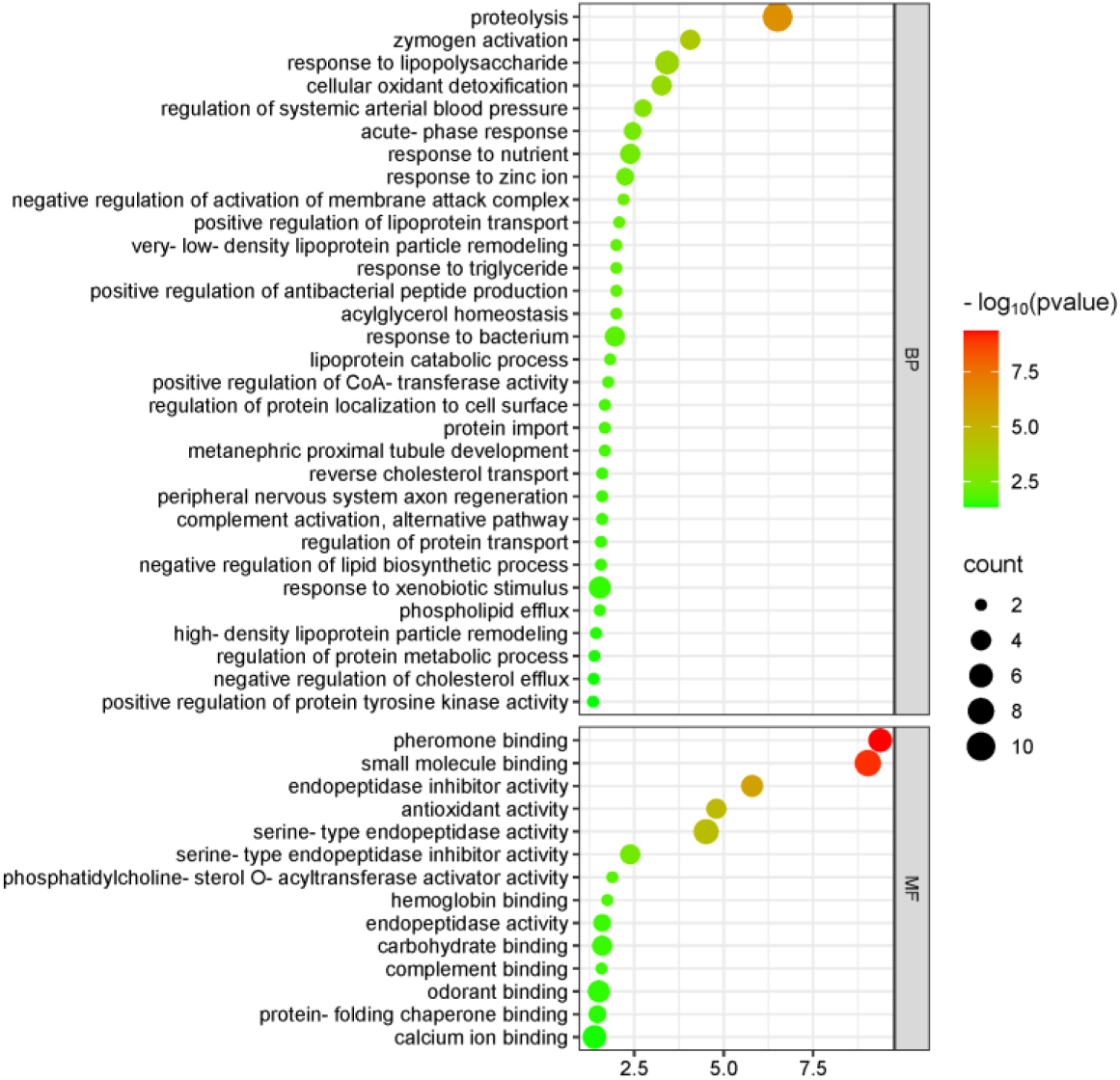
Enrichment analysis of biological processes and molecular functions of proteins containing differentially modified peptides from presence to absence or absence to presence.

KEGG enrichment analysis showed that significantly enriched pathways include renin-angiotensin system, complement and coagulation cascades, lysosome, and cholesterol metabolism (Figure 11).

**Figure 11.**
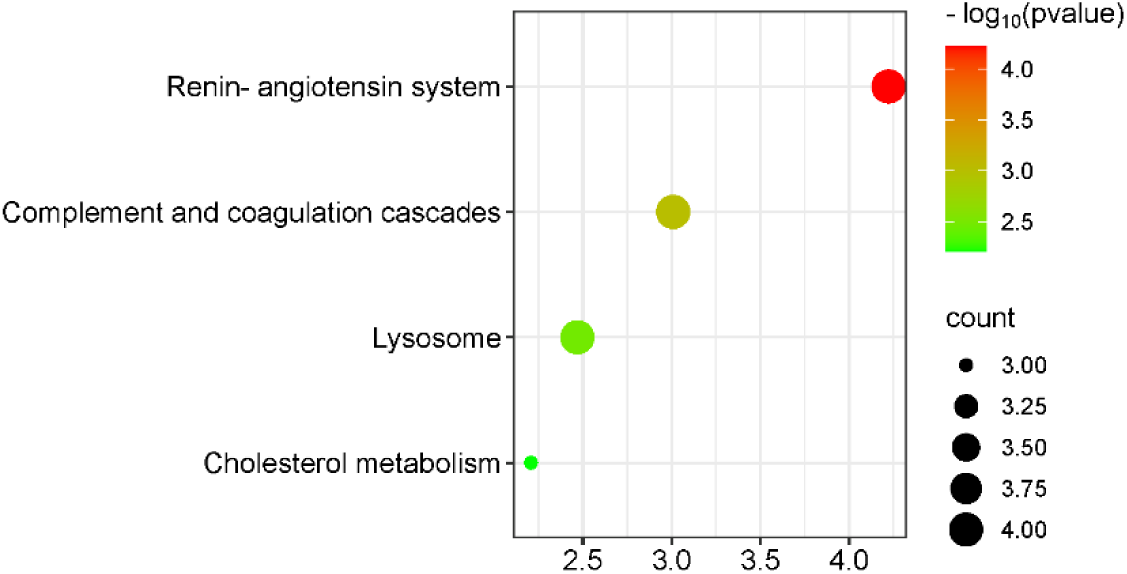
Enrichment analysis of the KEGG pathway of proteins containing differentially modified peptides from presence to absence or absence to presence.

## 4 Conclusion

The application of urine proteome helps to explore the known or unknown effects of drugs comprehensively and systematically, opening a new window to study the mechanisms of metformin.

---

**Supplementary Table 1.**
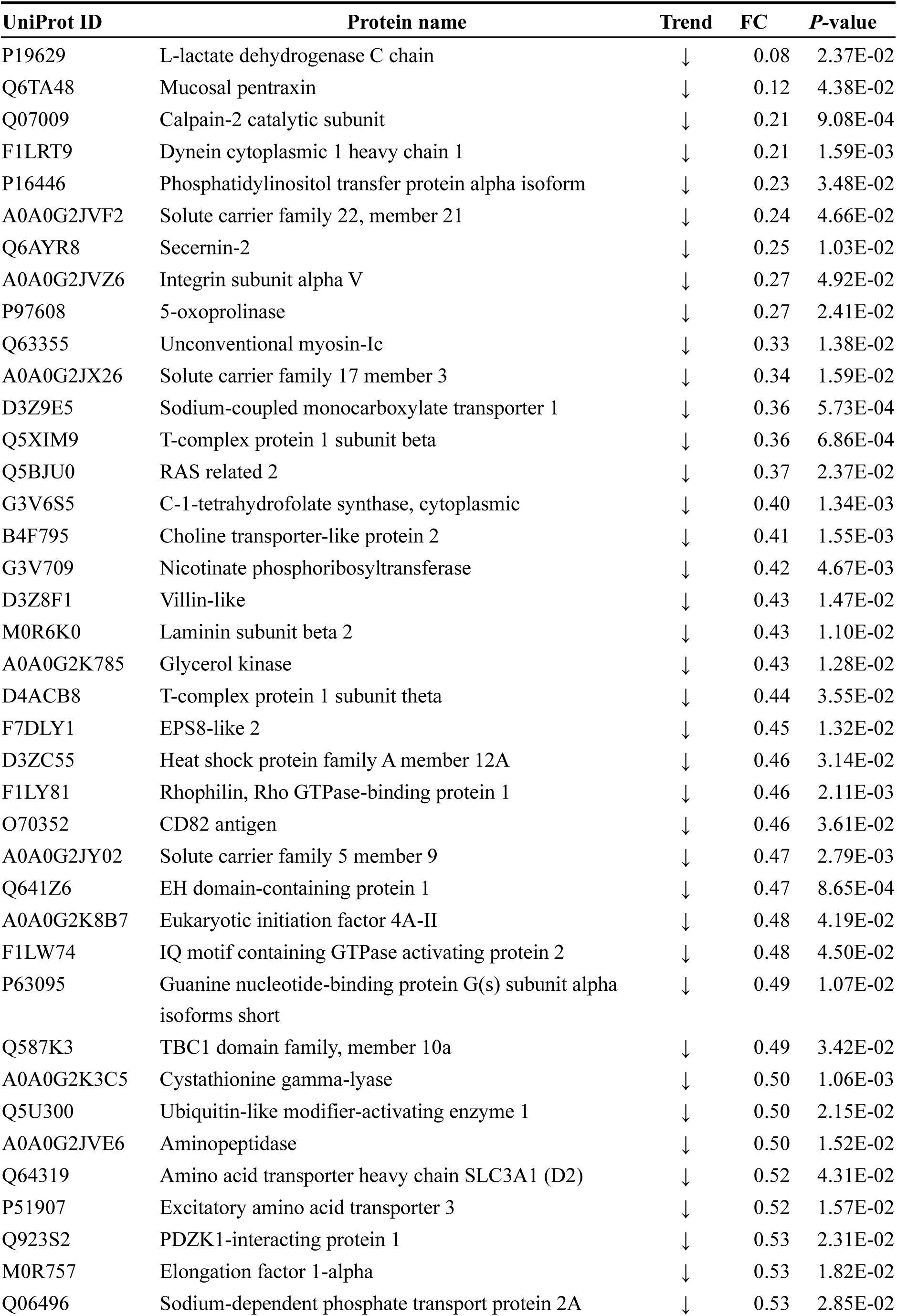
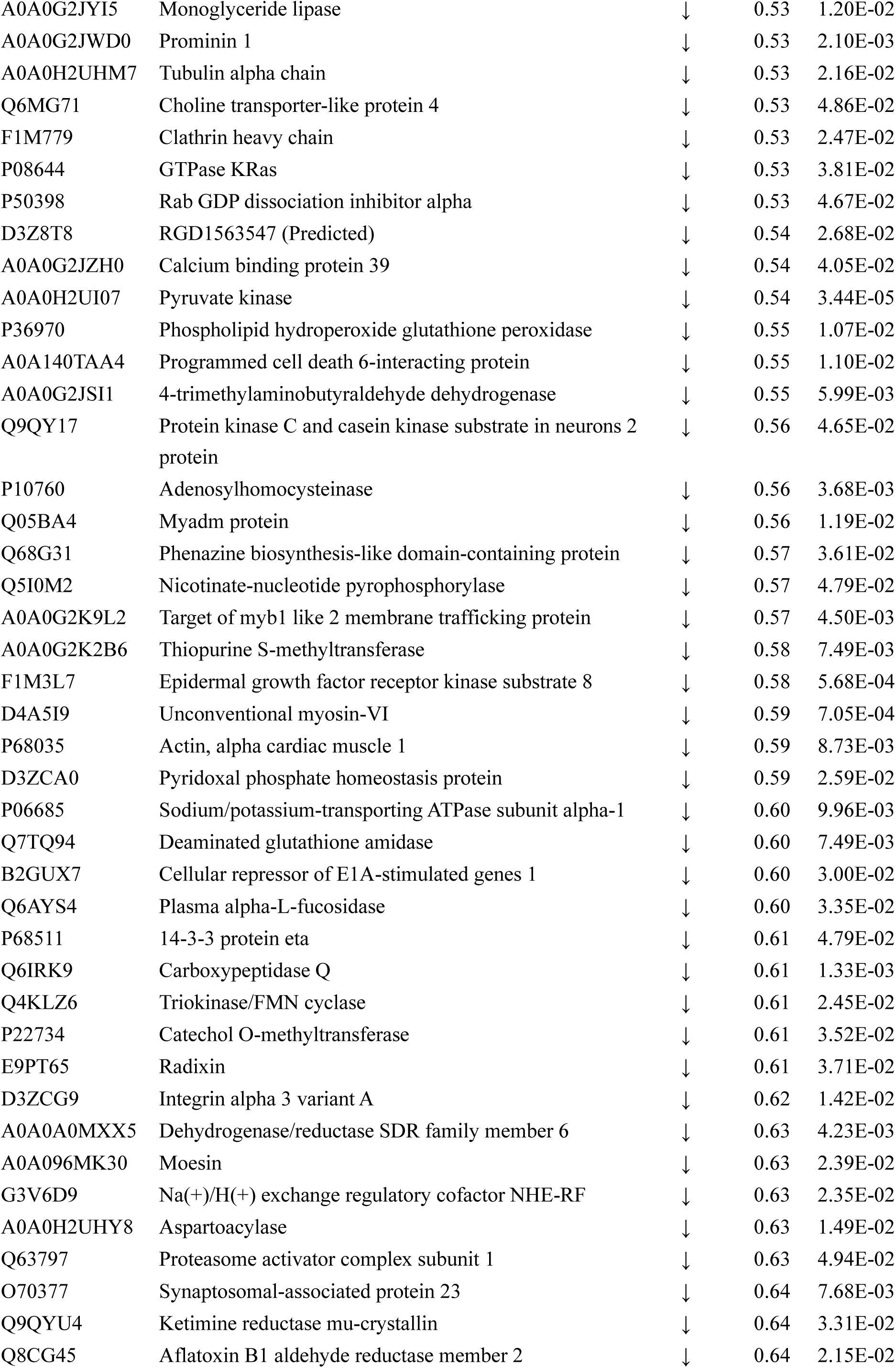
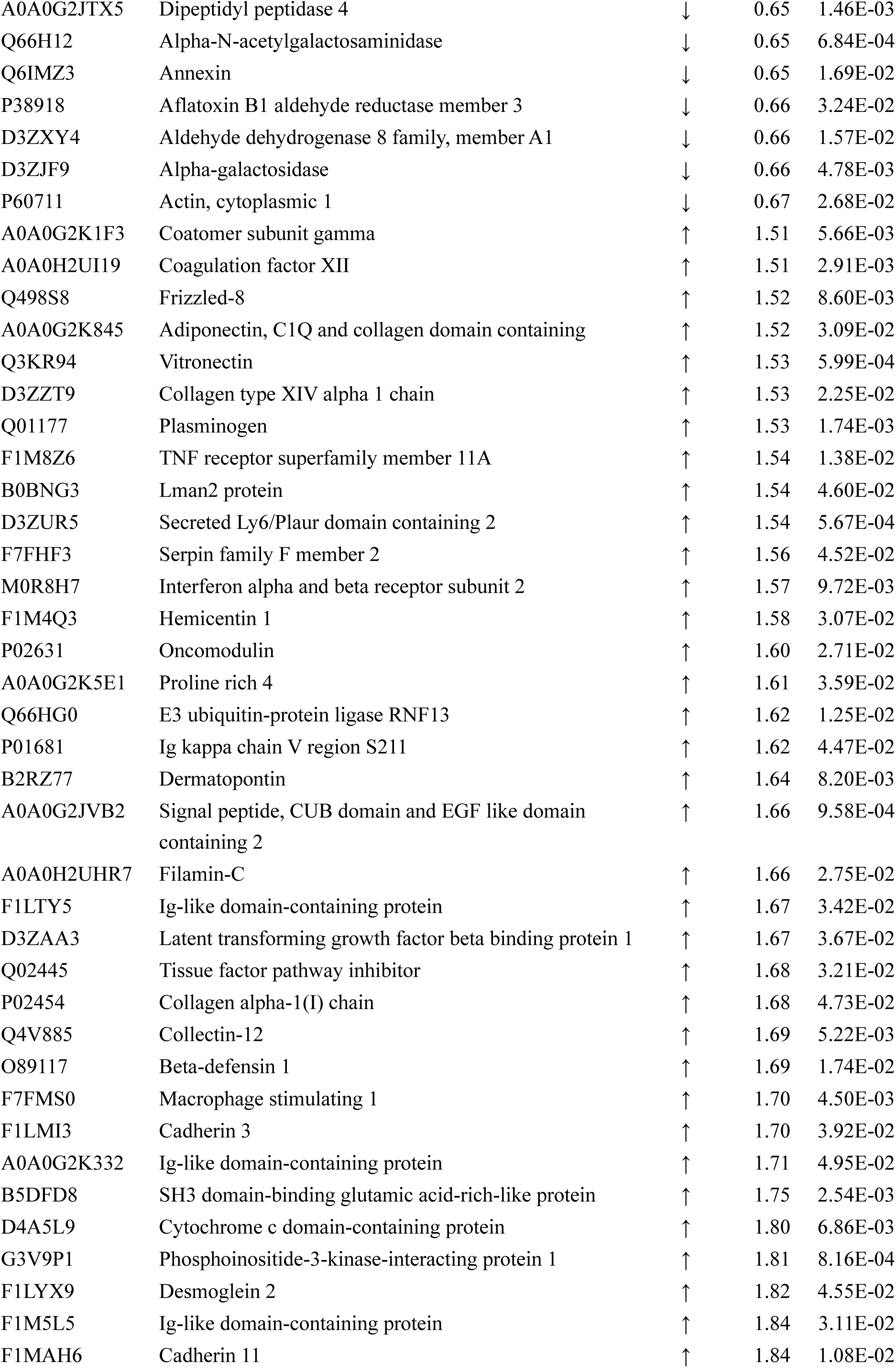
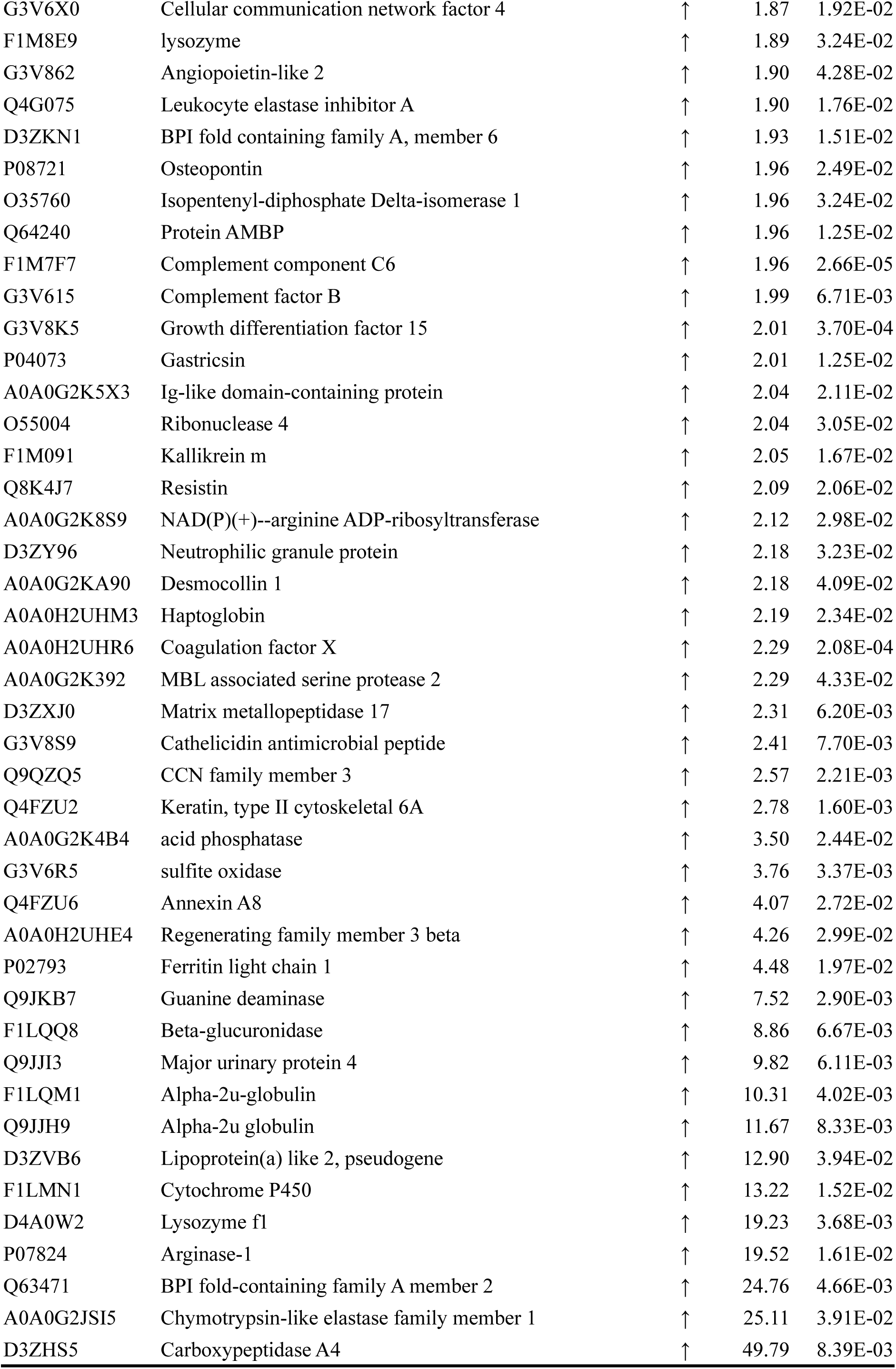
Differential proteins identified by comparison between the experimental group and the control group.

